# Importance of erythrocyte deformability for the alignment of malaria parasite upon invasion

**DOI:** 10.1101/611269

**Authors:** S. Hillringhaus, G. Gompper, D. A. Fedosov

## Abstract

Invasion of erythrocytes by merozoites is an essential step for the survival and progression of malaria parasites. In order to invade red blood cells (RBCs), parasites have to adhere with their apex to the RBC membrane. Since a random adhesion contact between the parasite and membrane would be too inefficient, it has been hypothesized that merozoites are able to actively re-orient toward apex-membrane alignment. This is supported by several experimental observations which show that merozoites frequently induce considerable membrane deformations before the invasion process. Even though a positive correlation between RBC membrane deformation and successful invasion is established, the role of RBC mechanics and its deformation in the alignment process remains elusive. Using a mechanically realistic model of a deformable RBC, we investigate numerically the importance of RBC deformability for merozoite alignment. Adhesion between the parasite and RBC membrane is modeled by an attractive potential which might be inhomogeneous, mimicking possible adhesion gradients at the surface of a parasite. Our results show that RBC membrane deformations are crucial for successful merozoite alignment, and require strengths comparable to adhesion forces measured experimentally. Adhesion gradients along the parasite body further improve its alignment. Finally, an increased membrane rigidity is found to result in poor merozoite alignment, which can be a possible reason for the reduction in the invasion of RBCs in several blood diseases associated with membrane stiffening.

**STATEMENT OF SIGNIFICANCE:** Plasmodium parasites invade erythrocytes during the progression of malaria. To start invasion, the parasites have to re-orient themselves such that their apex establishes a direct contact with erythrocyte membrane. The re-orientation (or alignment) process is often associated with strong membrane deformations, which are believed to be induced by the parasite and are positively correlated with its alignment. We employ a mechanically realistic erythrocyte model to investigate the interplay of membrane deformations and merozoite alignment during parasite adhesion to an erythrocyte. Our model clearly demonstrates that erythrocyte membrane deformations are a key component of successful parasite alignment, since the re-orientation of parasites at rigidified membranes is generally poor. Therefore, our results suggest a possible mechanism for the reduction in erythrocyte invasion in several blood diseases associated with membrane stiffening.

## INTRODUCTION

Malaria remains one of the most devastating diseases in the world, especially in African and South Asian regions, claiming over 400 000 lives per year (1). This motivates significant research efforts directed toward understanding various aspects and stages of malaria infection (2–4). Malaria is caused by a unicellular parasite from the genus *Plasmodium* which is transmitted to humans through a mosquito bite. Five different types of malaria parasites are known to infect humans. Among them, *Plasmodium falciparum* causes most severe cases of the infection. During the blood stage of malaria, merozoites invade red blood cells (RBCs) and asexually reproduce inside them. The invasion of RBCs by merozoites is a critical step in their survival (3–5), since inside RBCs the parasites remain invisible to the host’s immune system. As a result, this step in malaria has attracted considerable scientific interest, because it can reveal potential targets for antimalarial drugs (3).

Merozoites possess an egg-like shape with an average diameter of about 1 μm (6). Their apex contains all required machinery to invade RBCs (7). However, in order to start the invasion process, parasites have to establish first a direct contact between their apex and RBC membrane (5, 7). For this purpose, merozoites have a surface coat with a number of embedded proteins which can bind to RBC membrane (7–9). The first contact between the merozoite and RBC can be considered to occur with a random orientation. However, such random parasite adhesion is unreliable for the establishment of the direct apex-RBC contact, because after about three minutes, parasites become non-viable and are not able to invade RBCs anymore (10). Therefore, it is hypothesized that merozoites are able to actively align their apex toward the RBC membrane (11). This alignment or pre-invasion stage occurs within the range of 2 s to 50 s (11–13), which is fast enough to proceed to RBC invasion afterwards.

Even though the alignment process has been observed in a number of experiments (12–15), the mechanisms that lead to a successful parasite alignment are still under discussion. One proposition is that the parasite re-orientation is guided by a gradient of adhesive agonists along the parasite’s body, such that their density increases toward the apex (6). This proposition is based on some evidence for the release of adhesive agonists from the parasite’s apex during invasion (3, 16, 17). An interesting feature which is frequently observed in the pre-invasion stage is RBC membrane deformations of various intensity (12–15). In fact, a recent experimental study (18) has suggested that a positive correlation between the magnitude of membrane deformations and the efficiency of RBC invasion exists. Interestingly, such membrane deformations subside right after the alignment is achieved and the merozoite starts initiating cell invasion. This suggests that the parasite may actively trigger membrane deformation in order to facilitate alignment. It has been hypothesized that the parasite may mediate RBC membrane properties by changing local concentration of calcium (Ca^2+^) ions (11, 19). However, recent experiments (20) provide evidence against this hypothesis, because RBC deformations are also observed in the absence of Ca^2+^ and calcium release by the parasite starts only at the invasion stage.

At present, several questions regarding possible mechanisms for the parasite alignment remain unanswered. Do parasites actively induce RBC membrane deformations or do they result from passive parasite adhesion? Is an adhesion gradient along the parasite’s body required for successful alignment? How do membrane deformations aid parasite alignment? Are they necessary for a proper alignment? In order to address these questions, we perform simulations of parasite adhesion to a RBC membrane (21). In particular, we focus on the so-called passive compliance hypothesis (20) which assumes that observed membrane deformations simply result from the parasite adhesion to the RBC. Parasite-membrane adhesion is modeled by an attractive potential, whose local strengths are adapted to represent different adhesion intensities and gradients. Our results show that the parasite-RBC adhesion interactions produce membrane deformations of various intensity. Both the required interaction strength (10) and the deformation intensity match experimental observations (12–15, 18). In further agreement with experiments (18), we find that membrane deformations significantly aid parasite alignment, since the parasite becomes partially wrapped by the RBC membrane, making a contact between the apex and RBC much more likely. Furthermore, simulations of parasite adhesion to a rigidified RBC show poor parasite alignment, indicating that RBC deformation is a key aspect for the successful alignment of a merozoite. Simulations with an adhesion gradient along the parasite’s body confirm that such gradients facilitate better alignment. However, strong adhesion gradients lead to a well-controlled, directed and fast re-orientation of the parasite with a nearly perfect alignment, while in experiments (18), parasite re-orientation is more erratic and slow. This result suggests that merozoites should not have strong adhesion gradients at their surface. Finally, the quality of the presented adhesion model to explain merozoite alignment is discussed.

## METHODS & MODELS

To investigate adhesion interactions between a RBC and a parasite, we employ models of cells with membranes having bending and stretching elasticity, which are embedded into a fluid represented by the dissipative particle dynamics (DPD) method (22, 23).

### Red blood cell and parasite models

The RBC membrane is described by a triangulated network model with *N*_rbc_ vertices that are distributed at the membrane surface of the cell. These vertices are connected by *N*_*S*_ springs and form *N*_*T*_ triangles. Mechanical properties of the RBC are described by the potential energy

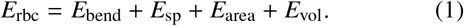

Here, the term *E*_bend_ represents the bending resistance of the lipid bilayer with a bending rigidity *κ*. *E*_sp_ models the elasticity of the spectrin network that is attached to the cytoplasmic side of the RBC membrane. Lastly, *E*_area_ and *E*_vol_ constrain the area and volume of RBC membrane, mimicking incompressibility of the lipid bilayer and cell’s cytosol, respectively. The elasticity of a RBC is characterized by a Young’s modulus *Y* and a shear modulus *μ*. This model has been verified to properly reproduce RBC mechanics (24, 25) and membrane fluctuations (26).

Similar to the RBC, the parasite is modeled by *N*_para_ vertices distributed on its surface, see Fig. 1 A. The egg-like shape and the size of a merozoite are approximated by (6)

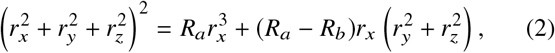

where *R*_*a*_ = 1 μm and *R*_*b*_ = 0.7 μm. The parasite is much less deformable than the RBC, as no deformations of parasite body are visible from different experimental observations (10, 18). Therefore, it is considered to be a rigid body, whose dynamics can be described by one equation for force and one equation for torque, which act on the parasite’s center of mass (27).

**Figure 1:**
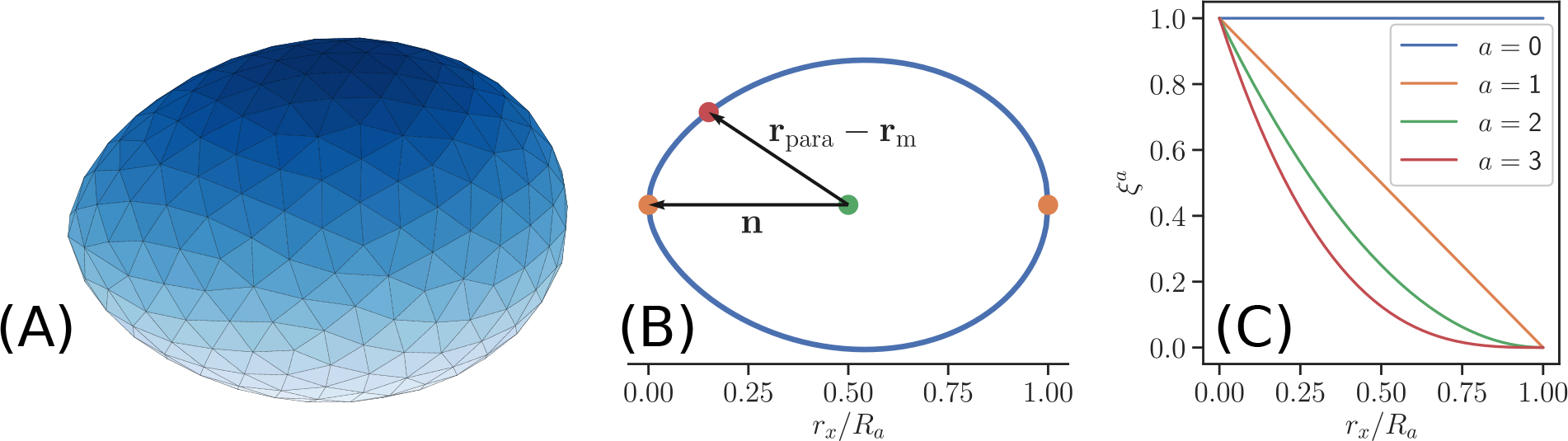
Parasite model and its adhesion interaction with RBC membrane. (A) The parasite is modeled by *N*_para_ vertices that form an egg-like shape given by Eq. 2. Motion of the parasite is modeled by rigid body dynamics. (B) The interaction for each parasite vertex depends on its relative position with respect to the parasite head (or apex), described by the dot product **n** · (**r**_para_ − **r**_m_) in Eq. 5. Here, 2**n** is a directional vector from the parasite back (*r*_*x*_/*R*_*a*_ = 1) to the head (*r*_*x*_/*R*_*a*_ = 0). (C) Function *ξ* describing a position-dependent density of adhesive proteins for various exponents *a*. *ξ* (**r**_para_) in Eq. 5 is chosen in such way that the interaction for *a* = 1 increases linearly along the parasite’s directional vector. Higher powers of *a* lead to interactions which are strongly localized around the parasite head.

### RBC-parasite adhesion interaction

The parasite and the RBC membrane interact through a Lennard-Jones (LJ) potential, which includes both repulsive and attractive parts. The pairwise interaction energy between one membrane vertex at **r**_rbc_ and one parasite vertex at **r**_para_ separated by the distance *r* = |**r**_para_ − **r**_rbc_| is given by

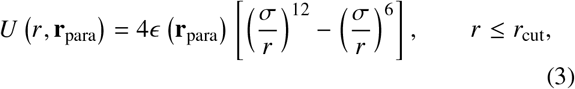

where *σ* is the characteristic length of repulsive interaction and *r*_cut_ is the cutoff radius for the adhesion potential. The potential is repulsive below 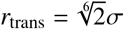 and attractive above. Note that the interaction strength *ϵ*(**r**_para_) may depend on a position r_para_ at the parasite surface. The total interaction energy is obtained by summing over all RBC-parasite vertex pairs.

The dependence of *ϵ* (**r**_para_) allows the control of interactions between the RBC membrane and parasite locally. For example, interaction gradients at the parasite surface can be introduced. To represent a gradient in adhesive agonists at the surface of the parasite, we define the surface density by

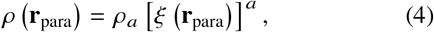

where *ρ*_*a*_ is the density coefficient and *a* is an exponent. The function *ξ* (**r**_para_) defines the distribution of agonists with respect to the position **r**_para_ at the parasite surface and is given by

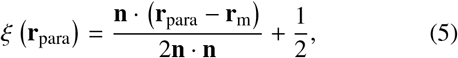

where **r**_m_ marks the mid point of the cylindrical symmetric axis. The vector **n**is the directional vector of the parasite, pointing from **r**_m_ towards its apex, see Fig. 1 B. For *a* = 1, the surface density of agonists decreases linearly with increasing distance in **n**-direction. Clearly, the exponent *a* can significantly shift the density of agonists toward the apex, as shown in Fig. 1 C.

In the discrete representation of the parasite in Fig. 1 A, each vertex corresponds to a small area *A*_c_ containing *A*_c_ *ρ* (**r**_para_) agonists. Under the assumption that the interaction strength for each agonist is *ϵ*_1_, we can define *ϵ* (**r**_para_) as

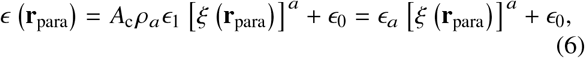

where *ϵ*_0_ represents an additional position-independent interaction between the RBC membrane and the parasite. In all simulations, *ϵ*_0_ is set to zero.

### Hydrodynamic Interactions

The RBC and parasite are embedded in a Newtonian fluid, which is modeled by a particle-based hydrodynamics method, dissipative particle dynamics (DPD) (22, 23). In short, the fluid environment is represented by a collection of fluid particles, which interact through three pairwise forces. Thus, the total force between particles *i* and *j* is given by

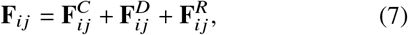

where 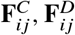, and 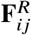 are conservative, dissipative, and random forces, respectively. The conservative force determines static pressure in the fluid and controls its compressibility. **F**^*D*^ governs fluid viscosity *η*, which depends on several simulation parameters and is evaluated using a reverse-Poiseuille flow setup (28, 29). Finally, **F**^*R*^ describes thermal fluctuations, such that the pair of dissipative and random forces serves as a thermostat, maintaining a desired equilibrium temperature. No-slip boundary conditions at the surface of the parasite and RBC membrane are enforced by an appropriate choice of the dissipative interaction between fluid particles and suspended cells (24).

### Simulation setup

The simulation setup consists of one RBC and a parasite suspended in a fluid. The simulated domain assumes periodic boundary conditions in all directions. The parasite is initially placed within the interaction range of the RBC, so that it can immediately adhere to the membrane. The initial orientation of the parasite is chosen such that the apex of the parasite is pointing away from the membrane, i.e. with its back to the RBC, the most unfavorable orientation for binding. Two different positions are investigated, parasite adhesion at the rim (or side) and at the highest point of the RBC membrane (see Fig. 2). For convenience, RBC center of mass is fixed by a harmonic spring, so that the cell does not diffuse away.

**Figure 2:**
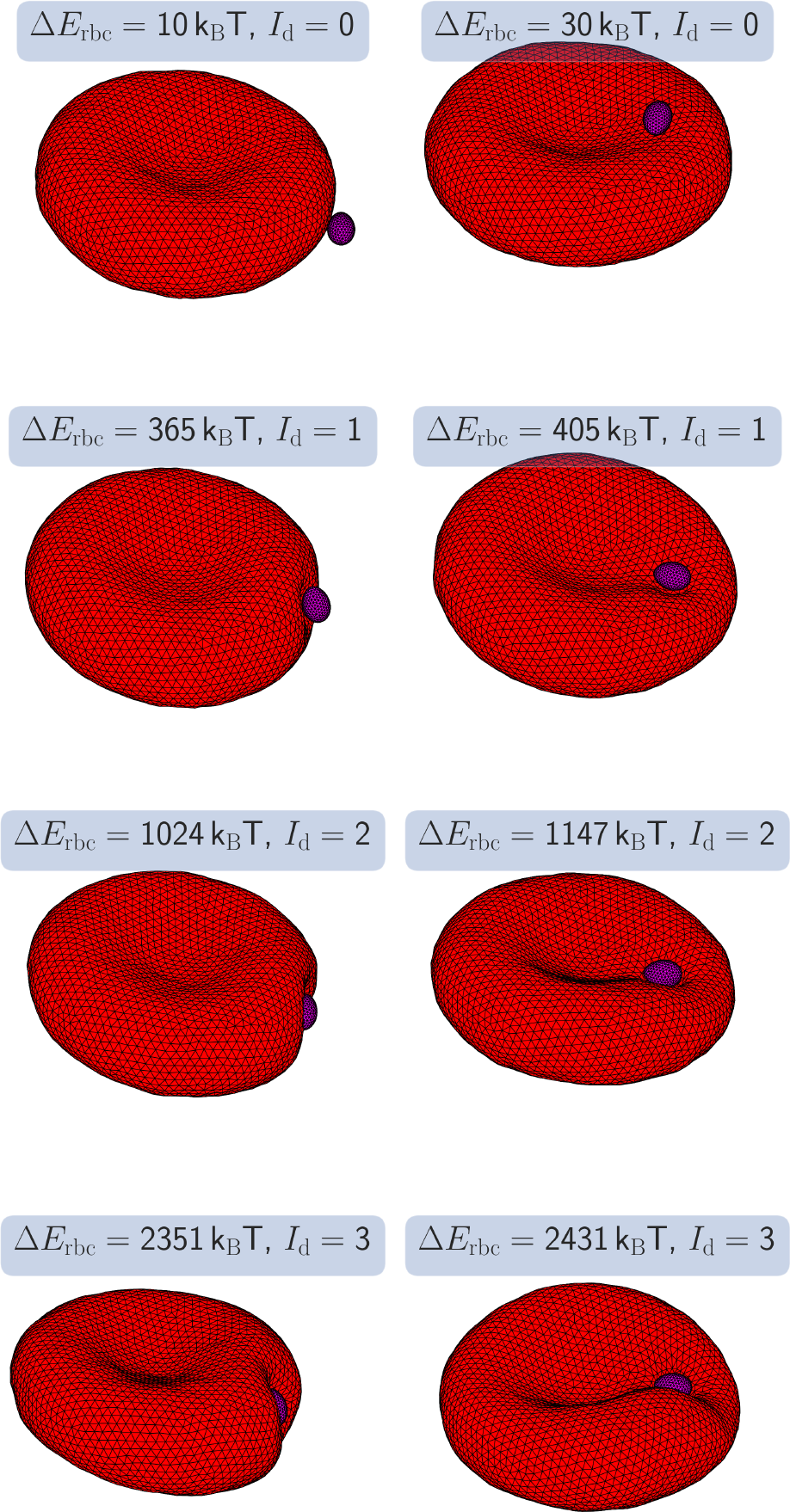
Snapshots of deformed RBCs due to parasite adhesion, including side (left column) and top (right column) contacts. Depending on the interaction strength the parasite induces membrane deformations of various intensity. These deformations are classified visually by a deformation index *I*_*d*_(see Table 2) and quantified by the deformation energy *E*_rbc_ shown in Fig. 3. Due to the interaction strength and partial wrapping of the parasite by the membrane, the parasite re-orients itself toward a configuration with a minimum total energy and shows no significant motion afterwards, see Movies S1-S2. Here, *a* = 0 (homogeneous adhesion).

Table 1. summarizes the main simulation parameters. To relate parameters in simulation and physical units, we define basic length, energy, and time scales. The length scale is defined by an effective RBC diameter 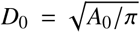, where *A*_0_ is the RBC membrane area. The basic energy scale is *k*_*B*_*T*, where *k*_*B*_ is the Boltzmann constant and *T* is temperature. Finally, the time scale *τ* corresponds to a RBC relaxation time defined as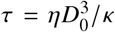. For average properties of a healthy RBC with *D*_0_ = 6.5 μm and *κ* = 3 × 10^−19^ J and for the fluid viscosity *η* = 1 mPa s, we obtain *τ* ≈ 0.92 s. All simulations were performed on the supercomputer JURECA (30) at Forschungszentrum Jülich. To obtain reliable averages of quantities of interest, all presented data points are averaged over about 10 statistically independent simulations.

**Table 1:** Overview of main parameters in simulation and physical units. The effective RBC diameter 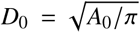 with *A*_0_ being the membrane area, the thermal energy *k*_B_*T*, and the characteristic RBC relaxation time 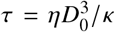 are selected as length, energy, and time scales. *κ* is the membrane bending rigidity, *Y* is the Young’s modulus, and *η* is the fluid’s dynamic viscosity. The number of vertices for the parasite *N*_para_ and RBC *N*_rbc_ are kept same in all simulations. *σ* and *r*_cut_ are parameters of the LJ potential. The RBC properties correspond to average characteristics of a healthy RBC.

| Parameter | Simulation value | Physical value |
| --- | --- | --- |
| $A_0$ | 133.5 | $133.5 \times 10^{-12} \text{ m}^2$ |
| $D_0$ | 6.5 | $6.5 \times 10^{-6} \text{ m}$ |
| $k_B T$ | 0.01 | $4.282 \times 10^{-21} \text{ J}$ |
| $\tau$ | 725.8 | 0.92 s |
| $\eta$ | $70 k_B T \tau / D_0^3$ | $1 \times 10^{-3} \text{ Pa s}$ |
| $\kappa$ | $70 k_B T$ | $3.0 \times 10^{-19} \text{ J}$ |
| $Y$ | $1.82 \times 10^5 k_B T / D_0^2$ | $1.89 \times 10^{-5} \text{ N m}^{-1}$ |
| $N_{\text{para}}$ | 310 | |
| $N_{\text{rbc}}$ | 3000 | |
| $\sigma$ | 0.15 | $0.15 \mu\text{m}$ |
| $r_{\text{cut}}$ | 0.4 | $0.4 \mu\text{m}$ |

## RESULTS

### Adhesion-induced membrane deformations

As a result of adhesion interactions between the parasite and RBC, the membrane can strongly deform (see Fig. 2 and Movies S1-S2), depending on the interaction strength. RBC deformation is quantified by the deformation energy Δ*E*_rbc_ defined as

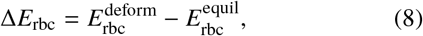

where 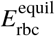 is the membrane energy for a biconcave RBC shape in equilibrium. Figure 3 presents membrane deformation energy as a function of the interaction strength *ϵ*_*a*_ for different exponents *a*. The deformation energy increases with increasing *ϵ*_*a*_ and strong deformations are induced by a partial wrapping of the parasite by the membrane in order to maximize the area of contact. Note that the adhesion between the parasite and RBC membrane results in a stable configuration after a very fast re-orientation, which does not change over time. The deformation energies are computed after this stationary configuration is reached. The data in Fig. 3 are for side contact. The results for top contact have a very similar dependence on *ϵ*_*a*_ and deviate by maximum 10 % from the values in Fig. 3. This is also consistent with the visual observations of RBC deformation in Fig. 2, showing membrane deformations of a similar degree for the both cases of RBC-parasite contact.

**Figure 3:**
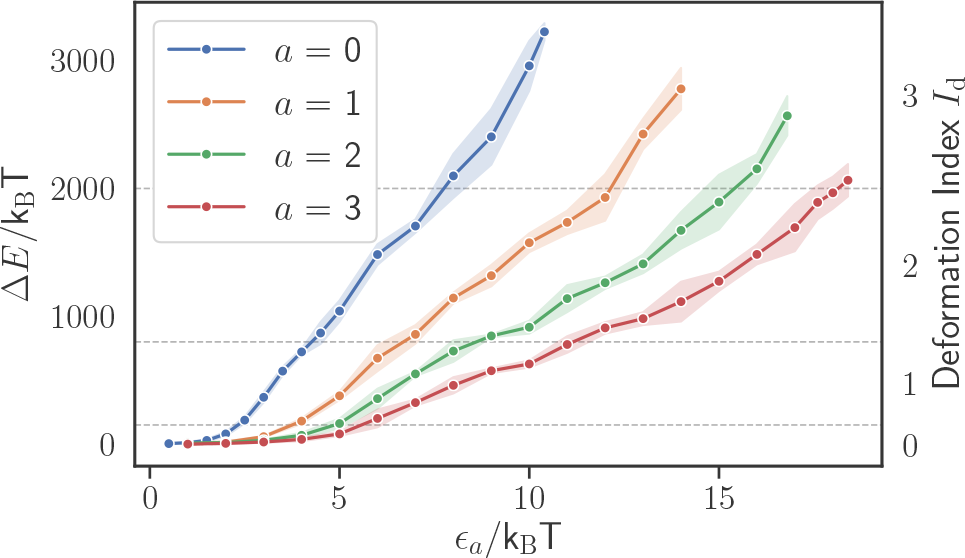
Deformation energy *E*_rbc_ and deformation index *I*_*d*_ as a function of the interaction strength *ϵ*_*a*_ for different exponents *a*. The shown values are for side contact between the parasite and RBC membrane. The lines for top contact are similar with a deviation up to 10 % from the shown results. The dashed lines indicate ranges of assigned deformation indices, see Table 2.

To characterize RBC deformations, a discrete deformation index *I*_*d*_ was introduced in experiments (18). This index divides membrane deformations into four categories and is assigned based on the visual inspection of RBC deformations. These categories are summarized in Table 2. In addition, we also associate the deformation indices with different ranges of deformation energy (see Table 2), using our simulation results. The RBC-parasite configurations in Fig. 2 are representative examples of different index categories.

**Table 2:** Definition of the deformation index *I*_*d*_, which is determined by a visual inspection of RBC membrane deformations (18). Connecting visual membrane deformations to the computed deformation energy Δ*E*_rbc_ allows us to define deformation energy ranges, which describe well different deformation indices.

| $I_d$ | Visible deformations | $\Delta E_{\text{rbc}}/k_B T$ |
| --- | --- | --- |
| 0 | No visible deformations | $[0, 150)$ |
| 1 | Small and local deformations | $[150, 800)$ |
| 2 | Partial wrapping of the parasite, local deformations. | $[800, 2000)$ |
| 3 | Wrapping of the parasite, RBC shape is globally deformed | $\geq 2000$ |

Δ*E*_rbc_ represents a deformation energy where all contributions are lumped together. Figure 4 shows different contributions to Δ*E*_rbc_ for the case of *a* = 0 as a function of the interaction strength *ϵ*_*a*_. The main contributions to the deformation energy correspond to bending elasticity of the lipid bilayer and shear elasticity of the spectrin network. Contributions from the area- and volume-conservation constraints are very small and can be neglected. For small membrane deformations (i.e. for *I*_*d*_ = 1), the bending energy contribution Δ*E*_bend_ dominates over Δ*E*_stretch_. For large RBC deformations with *I*_*d*_ = 2 or 3, the stretching energy contribution Δ*E*_stretch_ becomes dominant, pointing to significant stretching of the spectrin network. Note that for *a* > 0, the results for different contributions to Δ*E*_rbc_ are similar to the case *a* = 0.

**Figure 4:**
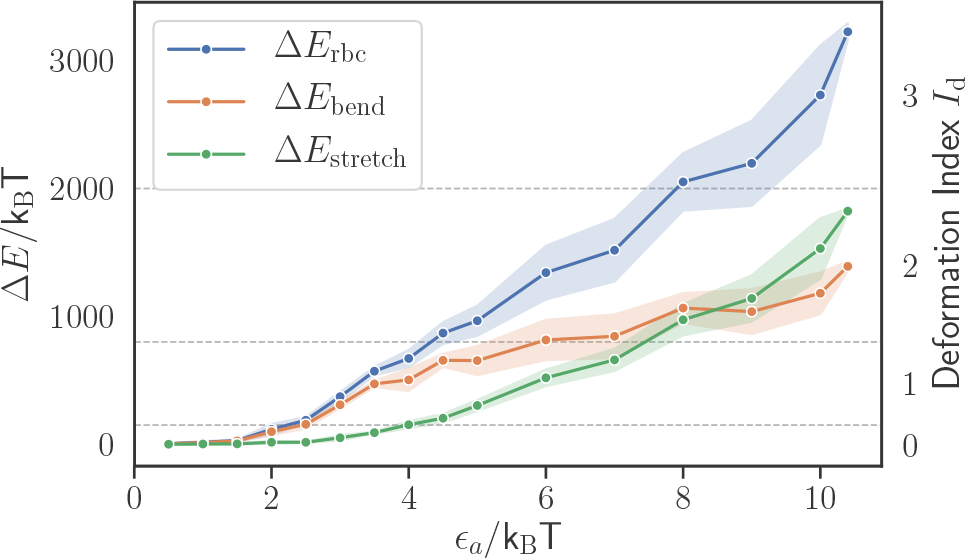
Different contributions to the deformation energy, including bending elasticity of the lipid bilayer and shear elasticity of the spectrin network, as a function of the interaction strength *ϵ*_*a*_. Small membrane deformations primarily correspond to membrane bending characterized by Δ*E*_bend_, while at large deformations, the contribution of stretching energy Δ*E*_stretch_ dominates. Here, *a* = 0 (homogeneous adhesion).

### Parasite adhesion force

As shown above, strong enough adhesion interactions between parasite and RBC lead to strong membrane deformations, similar to experimental observations (18). Interaction strength of spent merozoites has been measured in experiments by attaching a merozoite to two RBCs with the parasite in the middle (10). This is possible because spent parasites have lost their ability to invade RBCs, but still adhere to them. The elongation of one RBC pulled away by optical tweezers is used to quantify the force required for rupturing RBC-parasite adhesion contact. Experimental detachment forces are in the range of 10 pN to 40 pN (10).

We perform simulations mimicking these experiments to quantify adhesion forces for different interaction models and strengths. The corresponding simulation setup is shown schematically in Fig. 5 A. A parasite is adhered with its head (or apex) to a RBC that is pulled away with a constant velocity **v** (see Movie S3). The second RBC in the experimental setup is replaced by a harmonic spring with a spring constant 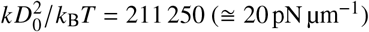, which tethers the parasite’s center of mass to its initial position. The pulling velocity is applied to a membrane rim position opposite to the parasite and is chosen such that the strain rate for RBC deformation remains close to 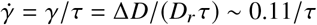 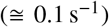, where *D*_*r*_ is the diameter of a RBC at rest and Δ*D* is the elongation of this diameter as a result of the applied strain *γ* (see Fig. 5 B). Detachment force is then measured as a maximum force *F*_ad_ = *k*Δ*L*_max_ on the harmonic spring tethering the parasite, as shown in Fig. 5 B. Here, Δ*L*_max_ is the spring elongation at the time when the connection between the parasite and RBC ruptures.

**Figure 5:**
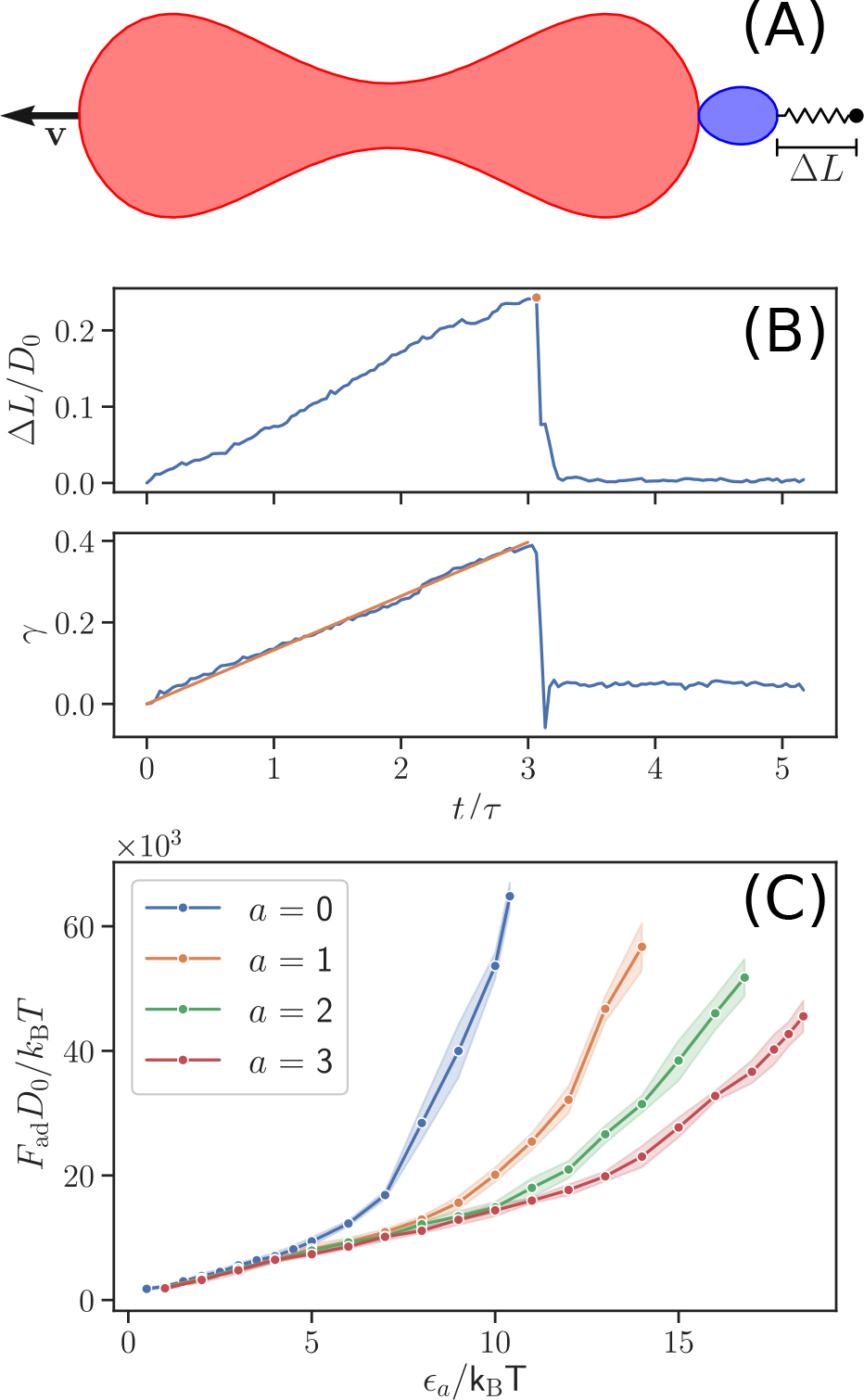
(A) Schematic illustration of a simulation to determine the detachment force *F*_ad_. A parasite, whose center of mass is tethered by a harmonic spring, is adhered to a RBC. The RBC is pulled away with a constant velocity **v** at a rim position opposite to the parasite (see Movie S3). (B) *F*_ad_ is measured through the elongation Δ*L* of the harmonic spring tethering the parasite, when it detaches from the RBC, i.e. the maximum measured force. Applied strain *γ* for RBC deformation corresponds to a nearly constant strain rate of 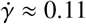. (C) *F*_ad_ as a function of the interaction strength *ϵ*_*a*_ for different values of *a*. The adhesion force increases for all values of *a* with increasing the interaction strength.

Figure 5 C presents detachment forces as a function of the interaction strength *ϵ*_*a*_ for different values of the exponent *a*. *F*_ad_ increases non-linearly with increasing *ϵ_a_*, because larger interaction strengths lead to a stronger wrapping (i.e. a larger interaction area) of the parasite by RBC membrane. The curve for *a* = 0 represents the steepest increase in *F*_ad_, as it corresponds to the strongest interaction between the parasite and RBC (see Fig. 1 C). The detachment forces from simulations in Fig. 5 C can be compared to experimentally measured forces (10), e.g. *F*_ad_ *D*_0_/*k*_*B*_*T* = 60×10^3^ corresponds to *F*_ad_ ≅ 40 pN. Therefore, *F*_ad_ in Fig. 1 C is smaller than 45 pN for all shown cases, so that the range of employed adhesion strengths realistically represents interactions between the parasite and the RBC membrane.

### Parasite alignment

For a successful RBC invasion, the parasite needs to align its apex (or head) toward cell membrane. Experiments indicate that the parasite head has to be in close proximity to the membrane surface and a successful invasion strongly correlates with a perpendicular alignment of the parasite toward RBC membrane (5, 8). Therefore, we introduce the head distance *d*_head_ and alignment angle *θ*, illustrated in Fig. 6, which allow the quantification of parasite alignment required for RBC invasion. *d*_head_ is defined as the distance between the parasite head **r**_head_ and a membrane vertex **r**_*i*_ that minimizes the distance *d*_head_ = min_*i*_ |**r**_head_ − **r**_*i*_|. *θ* is measured between the directional vector **n** of the parasite and the normal vector **n**_*i*_ of a triangular face whose center is closest to the parasite head, as sketched in Fig. 6. With these definitions, an optimal alignment is achieved for small values of *d*_head_ and an alignment angle *θ* ∼ *π*. Note, that this optimal value of *θ* may not be reached even for a perfect perpendicular alignment, since only the closest triangle is used to calculate *θ* and this triangle may not lie directly in front of the apex. Therefore, every angle *θ* ≥ 0.8*π* is considered to correspond to good alignment.

**Figure 6:**
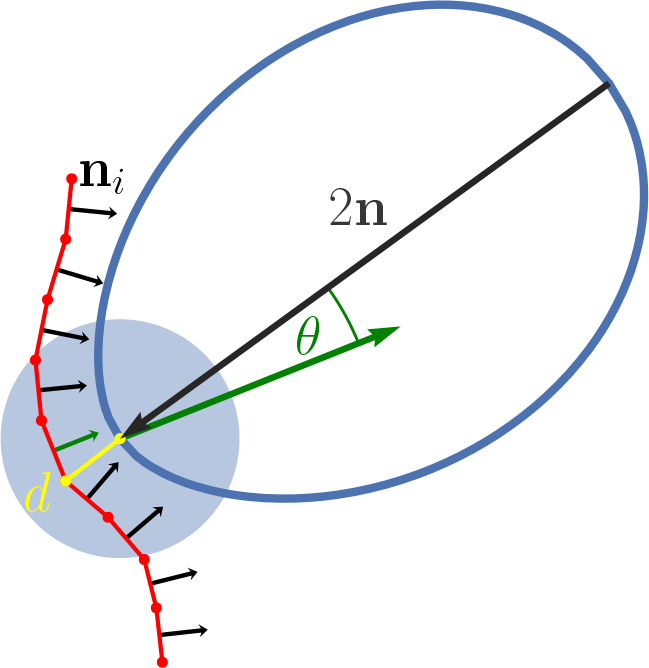
A sketch illustrating the measurement of a head distance *d*_head_ and alignment angle *θ*. *d*_head_ is calculated as the distance between the parasite head and the closest membrane vertex (yellow color). *θ* is determined as the angle between the directional vector **n** of the parasite and the normal of a triangular face (green arrow) whose center is closest to the parasite head. The alignment angle is calculated only if the head of the parasite is within a certain interaction range with the membrane, as indicated by the blue circle.

Figure 7 shows the two alignment characteristics as a function of the interaction strength *ϵ*_*a*_ for different exponents *a*. For all adhesion models, the values of *d*_head_ in Fig. 7 A decrease with increasing *ϵ*_*a*_ and closely approach the minimum possible distance represented by the repulsive range *σ* of the **LJ** interaction. The alignment angle *θ* in Fig. 7 B increases with increasing interaction strength. Thus, both characteristics show a positive correlation of parasite alignment with interaction strength *ϵ*_*a*_ or equivalently with RBC deformation. This is mainly due to the fact that a larger *ϵ*_*a*_ value leads to a stronger wrapping of the parasite by the RBC membrane, bringing the parasite head closer to the membrane surface. In addition, an interaction gradient along the parasite’s body for *a* > 0 aids the alignment, since such gradients favor parasite adhesion with an orientation of its head toward RBC membrane. The effect of interaction gradient on the parasite alignment can be clearly seen in Fig. 7, where both alignment properties are better for *a* > 0 in comparison to the case of *a* = 0. For instance, values of *θ* for *a* = 0 in Fig. 7 B do not closely approach π even for large *ϵ*_*a*_ values. Nevertheless, all models show good parasite alignment properties for interaction strengths *ϵ*_*a*_/*k*_*B*_*T* ≳ 5, which correspond to small levels of membrane deformations (*I*_*d*_ ≥ 1), as shown in Fig. 3.

**Figure 7:**
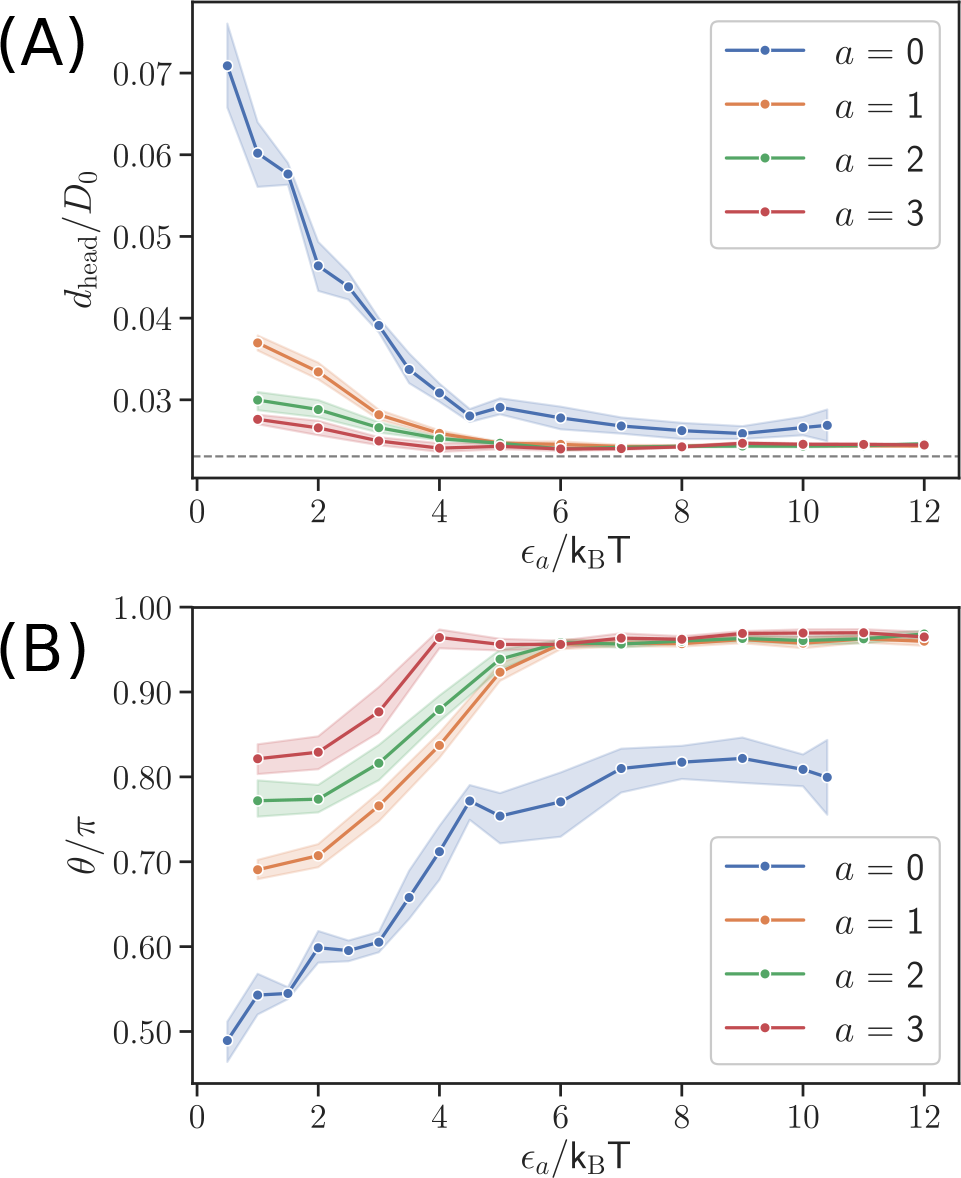
(A) Average head distance *d*_head_ and (B) average alignment angle *θ* for different interaction strengths *ϵ_a_* and exponents *a*. The dashed line in plot (A) represents the minimum possible distance *σ* of the repulsive LJ interaction. An optimal parasite alignment corresponds to small values of *d*_head_ and an alignment angle close to *π*. The both characteristics show a positive correlation between the parasite alignment and the interaction strength (or equivalently the level of RBC deformation).

Another important aspect of the parasite alignment is the average time required for this process. Experimental observations indicate that parasite alignment generally occurs within a time range between a few seconds and one minute (18). Figure 8 shows average alignment times for different interaction strengths *ϵ*_*a*_ and values of *a* measured in the simulations. The alignment time corresponds to a time difference between the moments the parasite starts to interact with the membrane and when it reaches its stationary adhesion configuration. The simulated alignment times in Fig. 8 are similar for all interaction models and are generally much smaller than 1.1 *τ* (≅1 s). This means that the alignment times in simulations are about two orders of magnitude smaller than those found experimentally (18).

**Figure 8:**
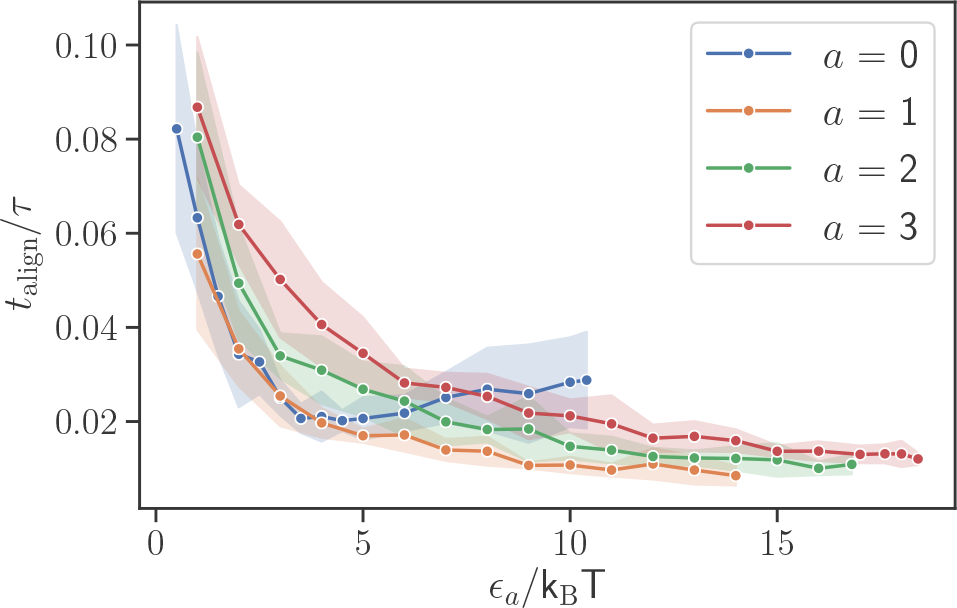
Average alignment time as a function of the interaction strength *ϵ_a_* for different exponents of *a*. All models result in alignment times *t*_align_/*τ* ≪ 1.

### Rigid RBC membrane

To investigate the effect of membrane deformations on the parasite alignment, simulations are performed with stiffened RBCs. Here, the bending rigidity and the Young’s modulus of RBC membrane are increased by two orders of magnitude in comparison to a healthy cell, so that the membrane can be considered rigid. As a result, the parasite does not induce deformations for all studied interaction strengths. Figure 9 shows the adhesion force *F*_ad_ as a function of *ϵ_a_* for *a* = 0 and *a* = 1. *F*_ad_ has a linear dependence on the interaction strength for both values of *a*. This is due to the fact that interaction area between the parasite and RBC is independent of *ϵ_a_* and remains constant, as the parasite cannot deform the RBC membrane. The detachment force can still reach magnitudes of up to 40 pN for large enough *ϵ_a_* values. *F*_ad_ is also similar for both interaction models.

**Figure 9:**
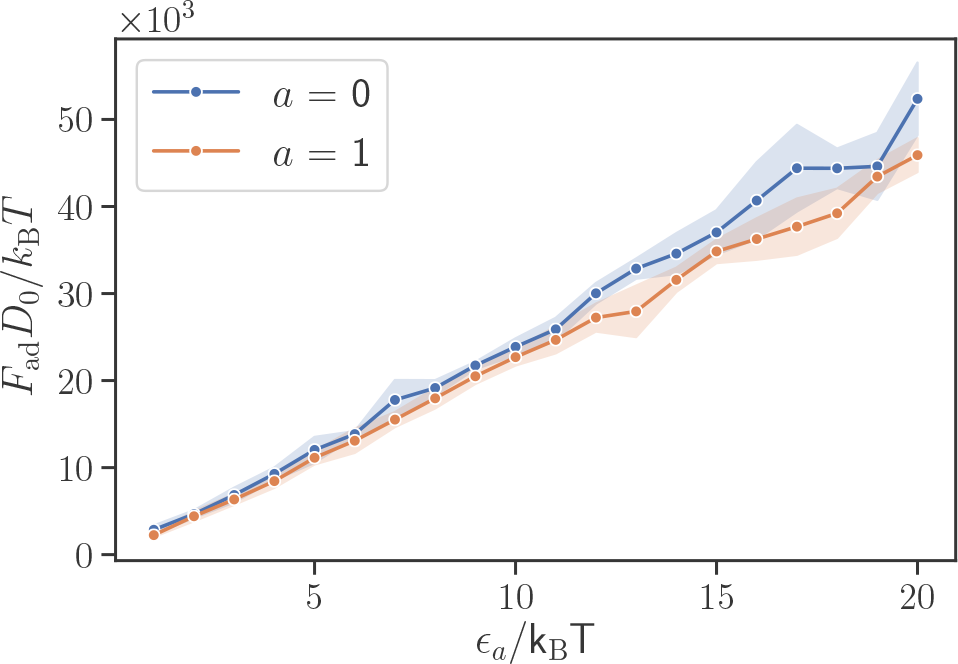
Adhesive force *F*_ad_ of the parasite to a rigid RBC for different interaction strengths *ϵ_a_* and values of *a*. *F*_ad_ for a rigid RBC grows slower with increasing *ϵ_a_* than that for a deformable RBC in Fig. 5 C, and has a nearly linear dependence on the interaction strength, as for rigid cell the adhesion area is independent of *ϵ_a_*.

Figure 10 presents the head distance *d*_head_ and the alignment angle *θ* for a parasite adhering to a rigid RBC. Both *d*_head_ and *θ* remain nearly constant independently of *ϵ_a_*. This means that the quality of parasite alignment is not influenced by the interaction strength and remains rather poor. This is due to the inability of the parasite to deform the RBC membrane, so that it positions itself sideways on the membrane (i.e. its directional vector is nearly perpendicular to membrane normal for *a* = 0, see Fig. 11, C), as this configuration represents a maximum adhesion energy. The case of *a* = 1 shows a better alignment in comparison to *a* = 0, since interaction gradient facilitates the re-orientation of parasite head toward the RBC membrane. However, both interaction models for parasite adhesion to a rigid RBC yield worse alignment results than those for deformable RBCs. This can be seen in Figure 11, which shows conformations of the parasite adhered to rigid and flexible membranes. For *a* = 1, the parasite has a good alignment for both membrane rigidities, since the interaction gradient (marked by color at the parasite surface) brings the parasite’s head close to the membrane (see Figs. 11 B, D). For *a* = 0, the energetically favorable adhesion configuration is a sideways positioning of the parasite with *θ* ≈ 0.5*π* (see Figs. 11 A, C and Movies S4-S5), which represents poor alignment. However, in case of a deformable RBC, the parasite can become partially wrapped by the RBC membrane, making the membrane-apex contact probable.

**Figure 10:**
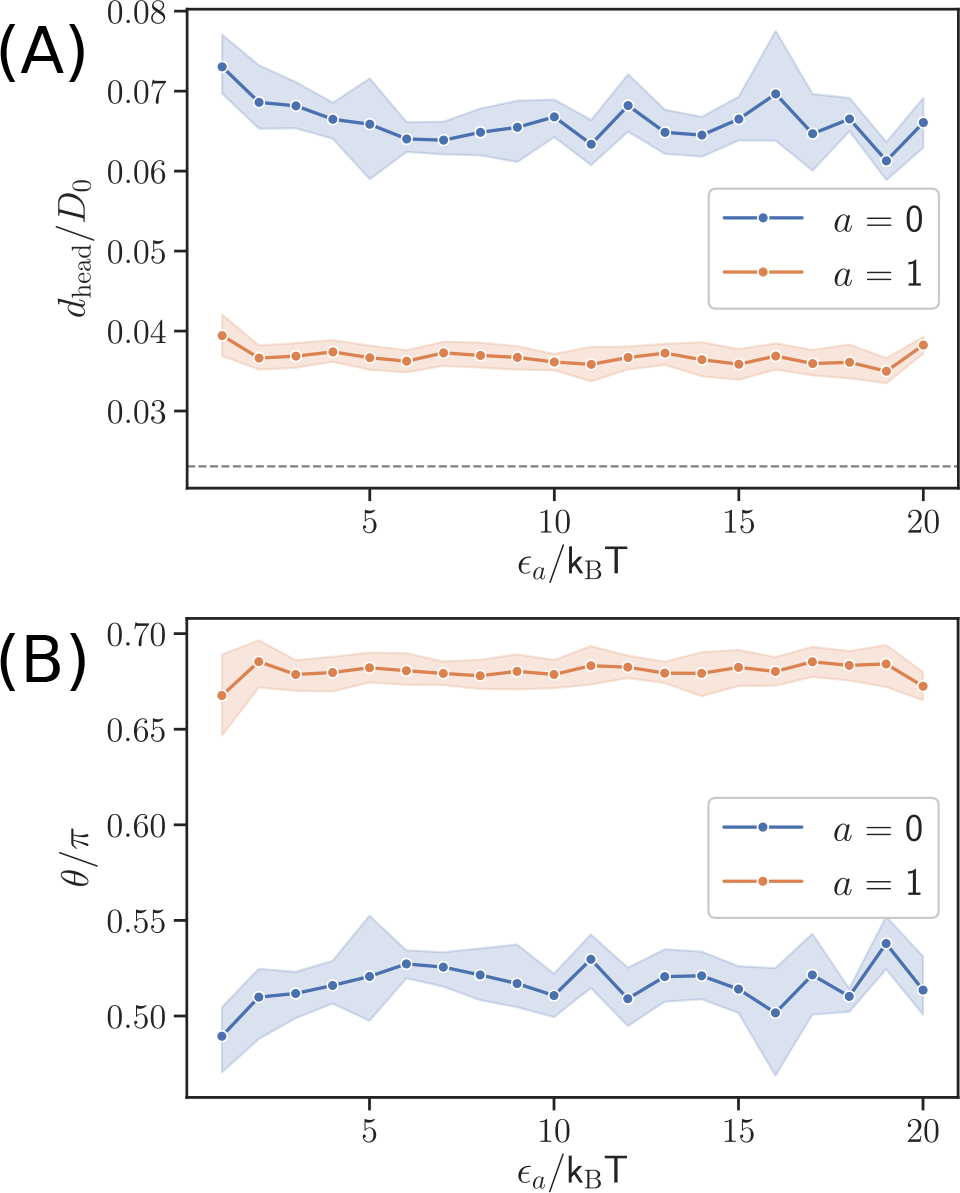
Alignment characteristics given by (A) head distance *d*_head_ and (B) alignment angle *θ* for the parasite interacting with a rigid RBC. Both characteristics are independent of the interaction strength *ϵ_a_* and generally show a poor parasite alignment.

**Figure 11:**
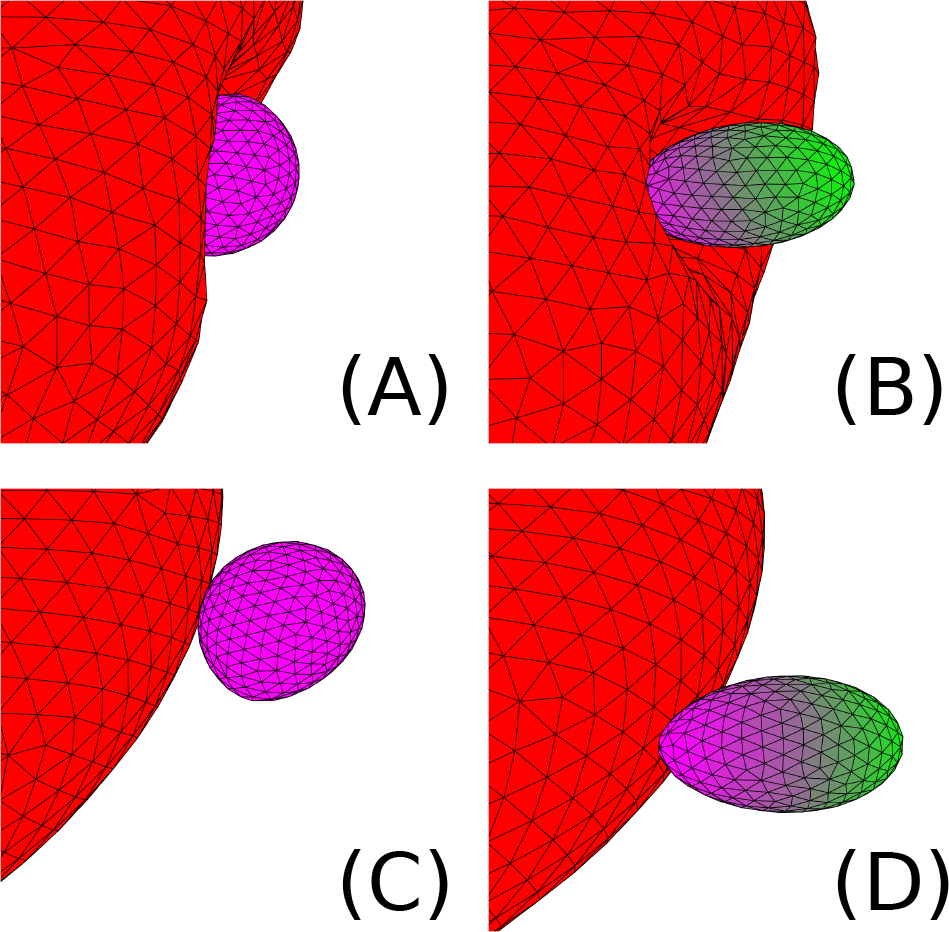
Comparison of parasite alignment at flexible RBCs (A, B) and rigid RBCs (C, D) for *a* = 0 (A, C) and *a* = 1 (B, D), where the interaction gradient is marked by a color code on the parasite surface (purple = maximal interaction strength, green = minimal interaction strength). For flexible RBCs, the observed alignment is better than for the rigid membrane, since the parasite can become partially wrapped by the flexible cell membrane (see Movies S4-S5). For *a* = 1, the alignment is good in both cases, since a configuration with the parasite’s head toward the membrane minimizes the total energy.

## DISCUSSION AND CONCLUSIONS

Our simulation results support the passive compliance hypothesis (20), which states that the alignment of merozoites arises from mechanical adhesion interactions between the parasite and RBC and induced membrane deformations. Here, both bending and stretching properties of the RBC membrane contribute to the resistance against parasite-induced deformations. For small local deformations, bending energy dominates, while strong membrane deformations lead to significant stretching of the RBC spectrin network. More importantly, the adhesion forces required for such deformations are within the range of experimentally measured forces 10 pN to 40 pN (10). The detachment force increases super-linearly with elevated interaction strength, because the parasite becomes strongly wrapped by RBC membrane, resulting in a significant increase of the interaction area. Thus, an increase in the interaction strength quickly leads to a very stable parasite-membrane adhesion. Both, the level of deformations and the adhesion forces well reproduce experimentally observed behavior (10, 18).

Analysis of parasite alignment characteristics such as the head distance *d*_head_ and the alignment angle *θ* shows that stronger membrane deformations lead to a better parasite alignment for all considered interaction models. The primary reason for the good alignment due to strong parasite-RBC interactions is that the parasite gets partially wrapped by RBC membrane, facilitating a contact between the membrane and the parasite head. For instance, in case of weak adhesion interactions, which do not induce significant membrane deformations, an equilibrium adhesion configuration corresponds to the parasite lying on its side due to the parasite’s egg-like geometry. This configuration makes a contact between the parasite head and the membrane unlikely. Therefore, strong membrane deformations serve as a hallmark of efficient parasite alignment followed by RBC invasion. This result compares favorably with recent experiments (18), where RBC deformations characterized by a deformation index were found to correlate positively with the parasite invasion frequency.

Furthermore, parasite models with an adhesion gradient along the parasite body show that such gradients facilitate a better alignment, as they introduce stronger adhesion interactions toward the parasite’s head. Strong enough adhesion gradients result in a perfect parasite alignment, which is generally not observed in experiments. In addition, the existence of adhesive gradients leads to very fast parasite alignment in a sub-second regime. The main reason for this quick reorientation due to adhesion gradients is that they lead to a well-controlled, directed and fast motion of the parasite toward perfect alignment by maximizing the interaction area. A comparison with experimental observations in Ref. (18) shows that the alignment in our simulations is about two orders of magnitude faster and that the real motion of the parasite at RBC membrane is often much more erratic than the parasite motion modeled with an interaction gradient. These results suggest that there should not be strong permanent adhesion gradients along the parasite body, since they lead to very fast re-orientation times and suppress diffusive parasite behavior.

The importance of RBC membrane deformations for the parasite alignment is further emphasized by the simulations of parasite adhesion to rigid RBCs. These simulations show that there is no correlation between the adhesion strength and the quality of parasite alignment. Furthermore, the alignment quality for the parasite interaction with a rigid RBC is quite poor in comparison to a deformable membrane. This is due to the fact that the minimum energy for parasite adhesion to a rigid surface corresponds to a configuration, where the parasite lies on its side because of its egg-like shape. This adhesion configuration is independent of the interaction strength and represents a poor alignment. Addition of the interaction gradient along the parasite’s body improves the parasite alignment on a rigid RBC, but does not make it perfect as in the case for a deformable membrane. An increased rigidity of the RBC membrane is relevant for several blood diseases and disorders such as sickle-cell anemia (31), thalassemia (32), and stomatocytosis (33). For example, the invasion efficiency of merozoites is reduced for sickle cell and thalassemic RBCs (34). Our simulation study suggests that a poor parasite alignment due to RBC membrane stiffening can contribute to the reduction in the invasion of RBCs by merozoites.

Even though the parasite adhesion model with a fixed interaction potential reproduces certain aspects of the parasite alignment, it does not capture frequently-observed erratic dynamics of a merozoite at a RBC membrane. The fixed interaction potential leads to a stationary adhesion configuration, where the parasite does not exhibit significant motion. This model captures the behavior of spent or inactive parasites, but it fails to describe parasites which are still fit for RBC invasion and show much more vivid dynamics. One possible reason for this model behavior can be the activity of the parasite agonists. The fixed interaction potential mimics a situation where the density of adhesive agonists at the parasite surface is large enough to allow an averaged description. A more realistic approach for the parasite adhesion would be an explicit representation of the discrete nature of binding agonists, which can be modeled by the formation and dissociation of discrete bonds between the parasite and the RBC membrane. Such a model is likely to lead to a more dynamic parasite adhesion to RBC membrane due to the stochasticity of discrete bindings, as the interaction depends on bond dynamics. Clearly, further investigation is needed to clarify the discrepancies in the adhesion dynamics between model predictions and experimental observations and to establish a reliable mechanism for the parasite alignment process.

## AUTHOR CONTRIBUTIONS

S. H. performed simulations and analyzed the computational results; G.G. and D.A.F. designed the research project; all authors interpreted the results and wrote the manuscript.

## ACKNOWLEDGMENTS

We would like to thank Virgilio L. Lew and Pietro Cicuta from the University of Cambridge for insightful and fruitful discussions. S.H. acknowledges support by the International Helmholtz Research School of Biophysics and Soft Matter (IHRS BioSoft). D.A.F. acknowledges funding by the Alexander von Humboldt Foundation. We also gratefully acknowledge the computing time granted through JARA-HPC on the supercomputer JURECA (30) at Forschungszentrum Jülich.

